# (Un)common space in infant neuroimaging studies: a systematic review of infant templates

**DOI:** 10.1101/2021.09.08.459462

**Authors:** Alexander J. Dufford, C. Alice Hahn, Hannah Peterson, Silvia Gini, Saloni Mehta, Alexis Alfano, Dustin Scheinost

## Abstract

In neuroimaging, spatial normalization is an important step that maps an individual’s brain onto a template brain permitting downstream statistical analyses. Yet, in infant neuroimaging, there remain several technical challenges that have prevented the establishment of a standardized template for spatial normalization. Thus, many different approaches are used in the literature. To quantify the popularity and variability of these approaches in infant neuroimaging studies, we performed a systematic review of infant MRI studies from 2000 to 2020. Here, we present results from 833 studies meeting inclusion criteria. Studies were classified into 1) processing data in single subject space, 2) using a predefined, or “off the shelf”, template, 3) creating a study specific template or 4) using a hybrid of these methods. We found that across the studies in the systematic review, single subject space was the most used (no common space). This was the most used common space for DWI and structural MRI studies while fMRI studies preferred off the shelf atlases. We found a pattern such that more recently published studies are more commonly using off the shelf atlases. When considering special populations, preterm studies most used single subject space while, when no special populations were being analyzed, an off the shelf template was most common. The most used off the shelf templates were the UNC Infant Atlases (26.1%). Using a systematic review of infant neuroimaging studies, we highlight a lack of an established “standard” template brain in these studies.

## Introduction

A critical preprocessing step for the analysis of magnetic resonance imaging (MRI) data is spatial normalization (Friston et al., 1995; Poldrack, Mumford, & Nichols, 2011). Spatial normalization is the process of bringing brain volumes that have been acquired in different individuals into a common neuroanatomical common (or reference) space (Crivello et al., 2002; Fox, Perlmutter, & Raichle, 1985; Poldrack et al., 2011) and is typically performed in analyses across all modalities: structural MRI (Ashburner & Friston, 2000), diffusion MRI (Jones et al., 2002), and functional MRI (Poldrack et al., 2011). Spatial normalization to a common space is often necessary for image statistics to be computed across participants (Friston, 1994; Gee, Alsop, & Aguirre, 1997), which assumes that across participants brain structures occupy the same standard anatomical space in a consistent manner (Fox, 1995; Toga & Thompson, 2001). Spatial normalization is highly dependent on the common space template used and results may not be directly comparable if different templates are used. (Laird et al., 2010; Lancaster et al., 2007; Rohlfing, Sullivan, & Pfefferbaum, 2009). With the goal of rigor and reproducibility in mind, it is crucial for common spaces to be standardized across fields of neuroimaging to compare across studies (Fox, 1995; Friston et al., 1995). For adult neuroimaging studies, two common spaces (along with a standard coordinate system, or stereotaxic space) have emerged as standard common spaces for spatial normalization: Talairach space (Talairach & Tournoux, 1988) and MNI space (Evans et al., 1993) (Collins, Neelin, Peters, & Evans, 1994), with MNI space now considered the “standard” (Laird et al., 2010).

In recent years, MRI has had increased utilization as a methodological tool to examine brain development in infancy (Eyre et al., 2020; Gilmore, Knickmeyer, & Gao, 2018; Howell et al., 2019; Li et al., 2019). However, there is no standard common space template for infant brain studies. The adult MNI atlas is not appropriate for infant studies primarily due to vast neuroanatomical differences between the adult and infant brain. (Gaillard, Grandin, & Xu, 2001) Studies have shown the use of the adult MNI atlas in the analysis of infant neuroimaging studies introduces significant biases (Kazemi, Moghaddam, Grebe, Gondry-Jouet, & Wallois, 2007). There also exists major challenges in the development of a standardized infant brain common space. For example, brain development during the first year of life is rapid and dynamic with specific anatomical patterns of development for different ages (Gilmore et al., 2007; Knickmeyer et al., 2008). Second, high quality neuroimaging data is difficult to acquire in infancy due to low spatial resolution, low tissue contrast, and high participant motion (Shi et al., 2011; Xue et al., 2007). Third, common space templates are typically constructed based upon a large sample of high-quality neuroimaging data, thus making a template difficult to construct in infants (Shi et al., 2011). With these existing challenges, approaches to spatial normalization for infant neuroimaging studies have been largely inconsistent (Li et al., 2019; Oishi, Chang, & Huang, 2019; Shi et al., 2011). While “standard” infant atlases have been proposed (Oishi et al., 2019; Shi et al., 2011), it is currently unclear which common space approaches are used most frequently in infant neuroimaging. In this systematic review, we conducted a comprehensive literature search, review of common spaces used by each study, and analysis of common spaces for infant neuroimaging studies published between the years 2000-2020. By conducting this systematic review, we sought to understand the current state of the infant neuroimaging field in terms of the popularity and variability in spatial normalization methodology and to assist in the field of infant neuroimaging to adopt a “standard” common space moving forward.

## Methods

### Objective

In this systematic review, we aimed to summarize the approaches to spatial normalization used in infant neuroimaging studies between the years 2000 and 2020. We only included original quantitative research studies in the systematic review.

### Eligibility criteria

Quantitative research studies were excluded from the systematic review if they were: (1) published before the year 2000 (2) written in languages other than English (3) animal studies, case reports, review articles, clinical/radiologist review, not MRI of the brain, and methodological manuscripts (3) fetal MRI studies or participants were older than 18 months chronological age (4) articles using only other imaging modalities other than MRI, (e.g. fNIRS, PET, EEG).

### Search procedure and studies identified

We conducted a search on PubMed for infant neuroimaging studies that fit the eligibility criteria. Literature was compiled on September 2-3rd, 2020 using the following search string: “infant MRI” and “neonatal MRI”, “neonatal ‘fmri’”, “toddler ‘fmri’”, “‘toddler fmri’”, “preterm fmri”, “neonate(s)”, “infant(s)”, “(((infant) OR (neonate)) Or (newborn)) AND ((fmri) OR (MRI) OR (DTI))”. The initial search resulted in 37,782 manuscripts. The authors conducted screening and eligibility assessment based upon the eligibility criteria previously described using the web-tool Rayyan (Ouzzani, Hammady, Fedorowicz, & Elmagarmid, 2016). After the screening procedure had identified a subset of manuscripts that fit the eligibility criteria (833), the full articles were reviewed for eligibility and coded into 4 categories based upon the common space utilized in the study: (1) single subject space (e.g., analyses conducted in native space or no common space was used) (2) a study specific common space such that the common space was generated using the data in the study (e.g., tract-based spatial statistics or TBSS option “-n”) (3) an “off the shelf” atlas was used as the common space (e.g., the UNC Neonate Atlas) or (4) a hybrid approach to common space was utilized (e.g., more than one common space within the same imaging modality or different common spaces were used for each imaging modality). The results from the screening procedure are shown in the PRISMA Consort Chart (see **Figure 1**). We examined the breakdowns of common space by imaging modality: diffusion-weighted imaging (DWI), structural MRI, or functional MRI (fMRI)/resting-state (rsfMRI). Further, we examined the distributions of imaging modalities by publication year, common space by year, common space by age, and common space by special population. For the studies that utilized off the shelf atlases, we examined the breakdown of which atlases were most used.

**Figure 1.**
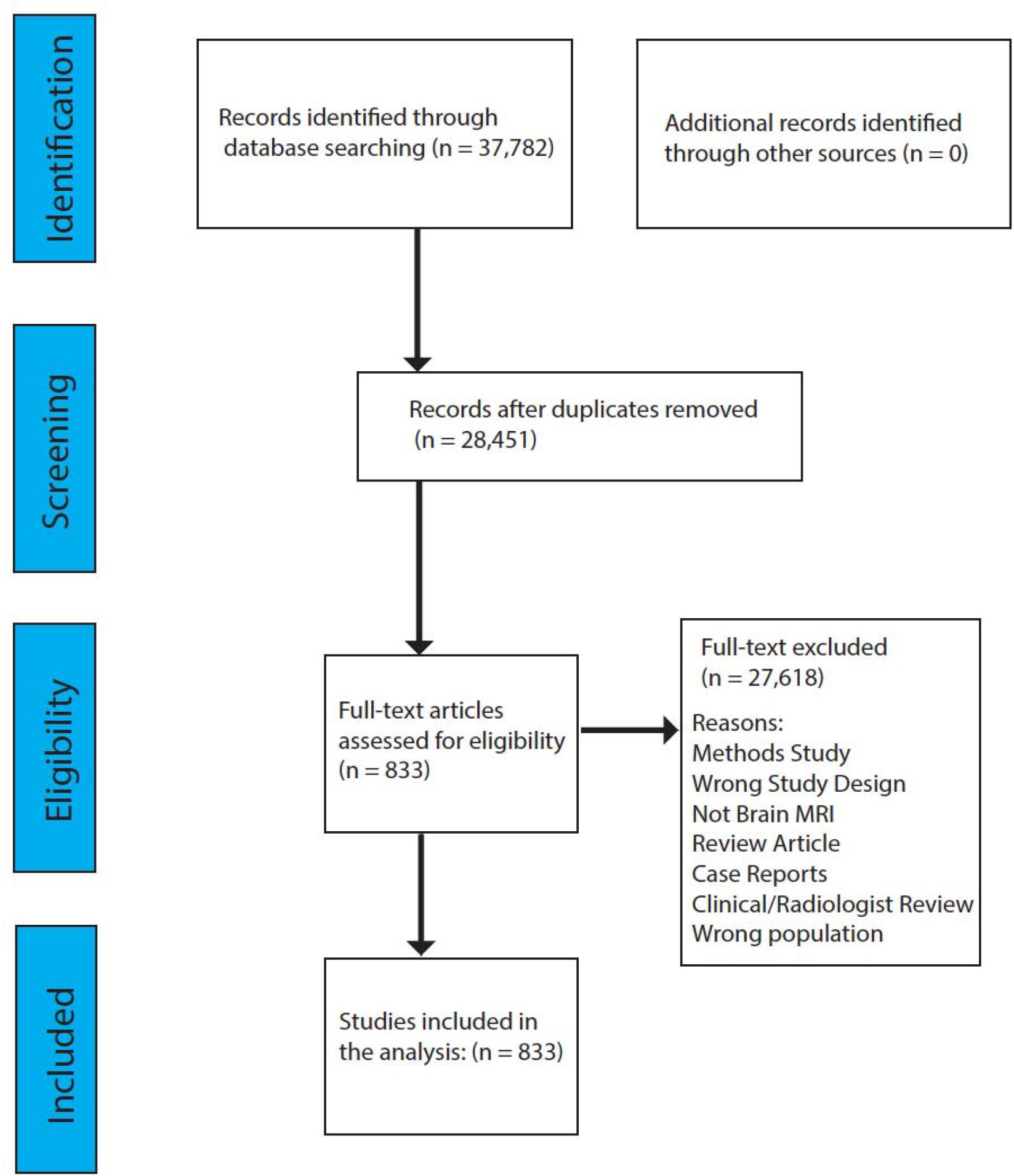
PRISMA diagram for the systematic review of infant common spaces.

### Statistical methods

For the analyses of common space (overall), by imaging modality, by age of the sample, by population, and by year, the distribution of “off the shelf” atlases, we used the count for each generated from the review of the full article. For a subset of the articles (n=298, articles from 2018-2020), we calculated the inter-rater reliability (Cohen’s kappa) of the agreement of the raters for classify articles into a common space category. For the calculation of Cohen’s kappa, we used the Kappa() function in the R package “irr” (Gamer, Lemon, Gamer, Robinson, & Kendall’s, 2012). To test for differences in distributions, we used a series of chisquare tests using the chisq.test() function in R. Additionally, we tested for a year published by modality interaction and a year published by population interaction using Poisson regressions (due to the count data). For the interaction testing, we first tested the dispersion of the data. If the data indicated the data was not overdispersed, a Poisson regression was conducted. To test the dispersion of the data, we used the dispersiontest() function from the R package “AER” (Kleiber, Zeileis, & Zeileis, 2020). Poisson regressions were conducted with the two predictors and interaction term e.g., Article Count ~ Common Space + Modality + CommonSpace*Modality.

### Data and code availability statement

The *R* script used for the analysis of the systematic review data is available at: (https://github.com/ajdneuro12/CommonSpace). The list of articles included in the analysis is available in the **Supplementary Information**.

## Results

### Common space by imaging modality

The analysis of the inter-rater reliability indicated “very good” agreement between raters: *K* = 0.945, *p* < 0.0001. According to our systematic review, the most used common space across imaging modalities was single subject space, followed by off the shelf atlases, study specific templates, and lastly by hybrid registrations (see **Figure 2**). We used Pearson’s chi-squared tests to examine if the distribution of the outcome (number of articles) was dependent on group (type of common space). Using three 3×2 contingency tables, we found evidence that the distribution of the article count was dependent on the common space when comparing single subject to off the shelf [*X^2^*(2, n = 671) = 103.75, *p* < .0001)], study specific to off the shelf [*X^2^*(2, n = 543) = 59.23, *p* < .0001)], single subject to off the shelf [*X^2^*(2, n = 598) = 19.08, *p* < .0001)]. When examined by the type of imaging modality, in DWI studies single subject space appeared to be the most prevalent (46.4% of all DTI studies), followed by study specific (31.4%), off the shelf (17.6%) and hybrid (4.6%). For structural MRI studies, single subject space and off the shelf atlases represented the majority (with 40.4% and 36.3% prevalence respectively), followed by registration to a study specific template (18.8%) and hybrid common space (4.5%). For fMRI and rsfMRI studies, off the shelf templates were the most common approach (60.1%), followed by study specific (20.9%). Analyses in single subject space (11.6%) and using hybrid methods (7.4%) were the least common for this imaging modality. Studies using a different common space registration method for each modality included in the paper were not included in the hybrid category but counted once per each modality and type of registration.

**Figure 2.**
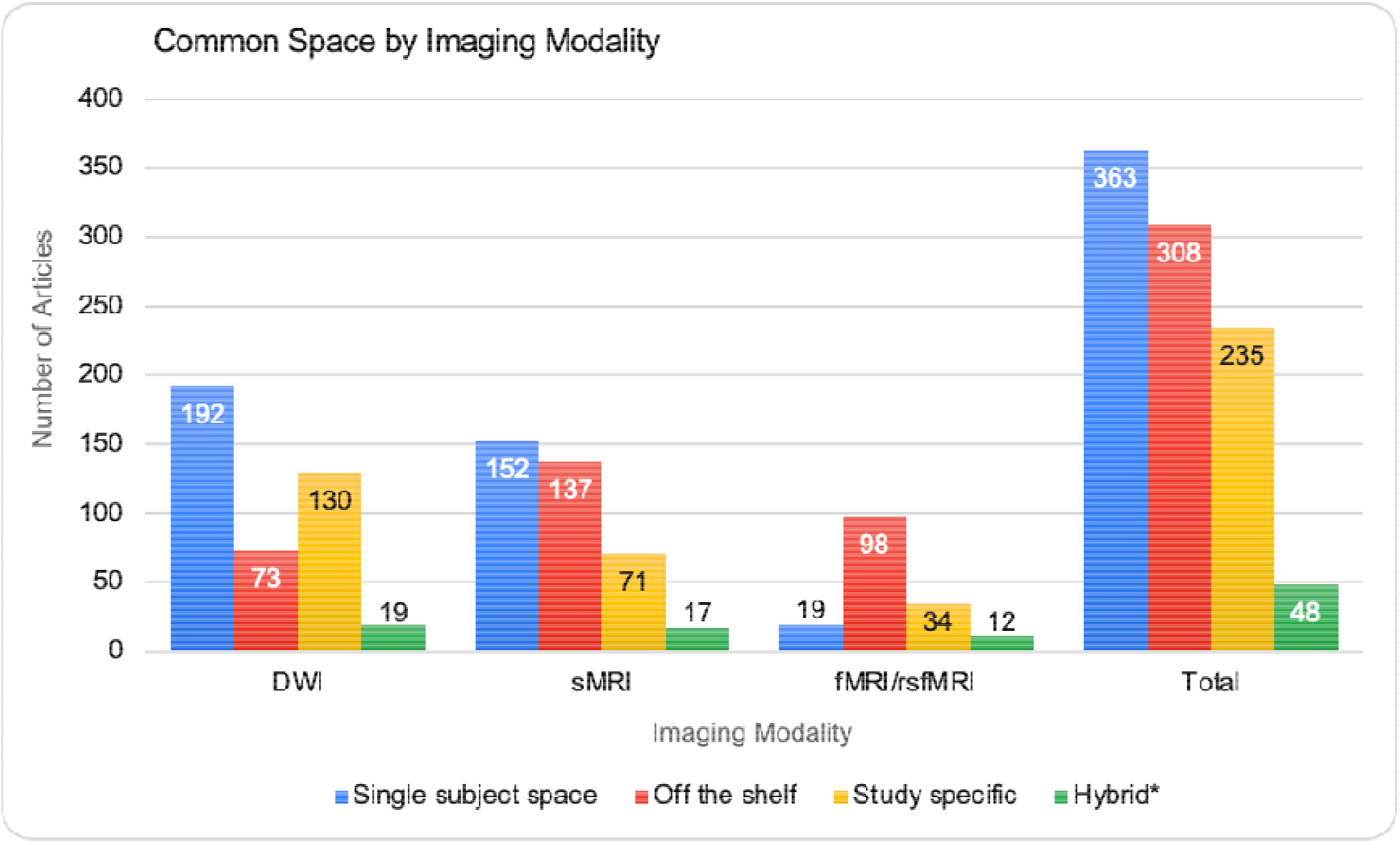
Frequency distribution of different common space registrations grouped by imaging modality and across all modalities (Total). The DWI (diffusion-weighted imaging) category includes studies employing diffusion tensor imaging (DTI), diffusion kurtosis imaging (DKI), and any other modality based upon diffusion weighted imaging (e.g., neurite orientation dispersion and density imaging). The sMRI category refers to structural MRI studies, and fMRI/rsfMRI to task and resting state functional MRI studies.

### Imaging modality by publication year

Results from the systematic review indicate that across imaging modalities, there was an increase in the number of studies utilizing an infant cohort from 2000 to 2020 (see **Figure 3**) with only a handful between 2000–2002 to over 350 between 2018–2020. In general, infant fMRI studies lag DWI and sMRI studies in terms of number of publications per year.

**Figure 3.**
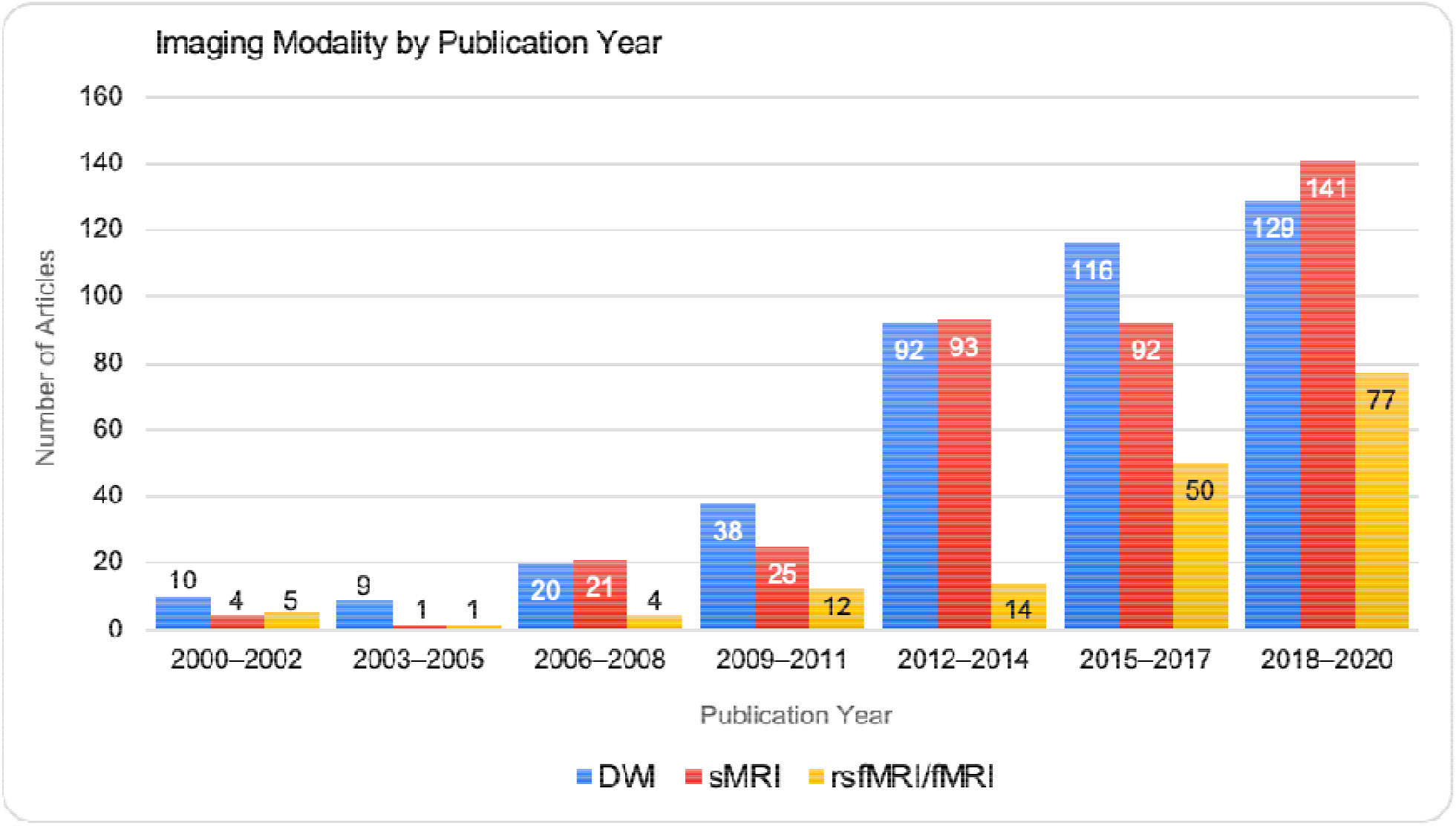
Frequency of imaging modalities across 20 years of infant MRI publications ranging from years 2000 to 2020. Distribution was analyzed in 3-year increments.

**Figure 4.**
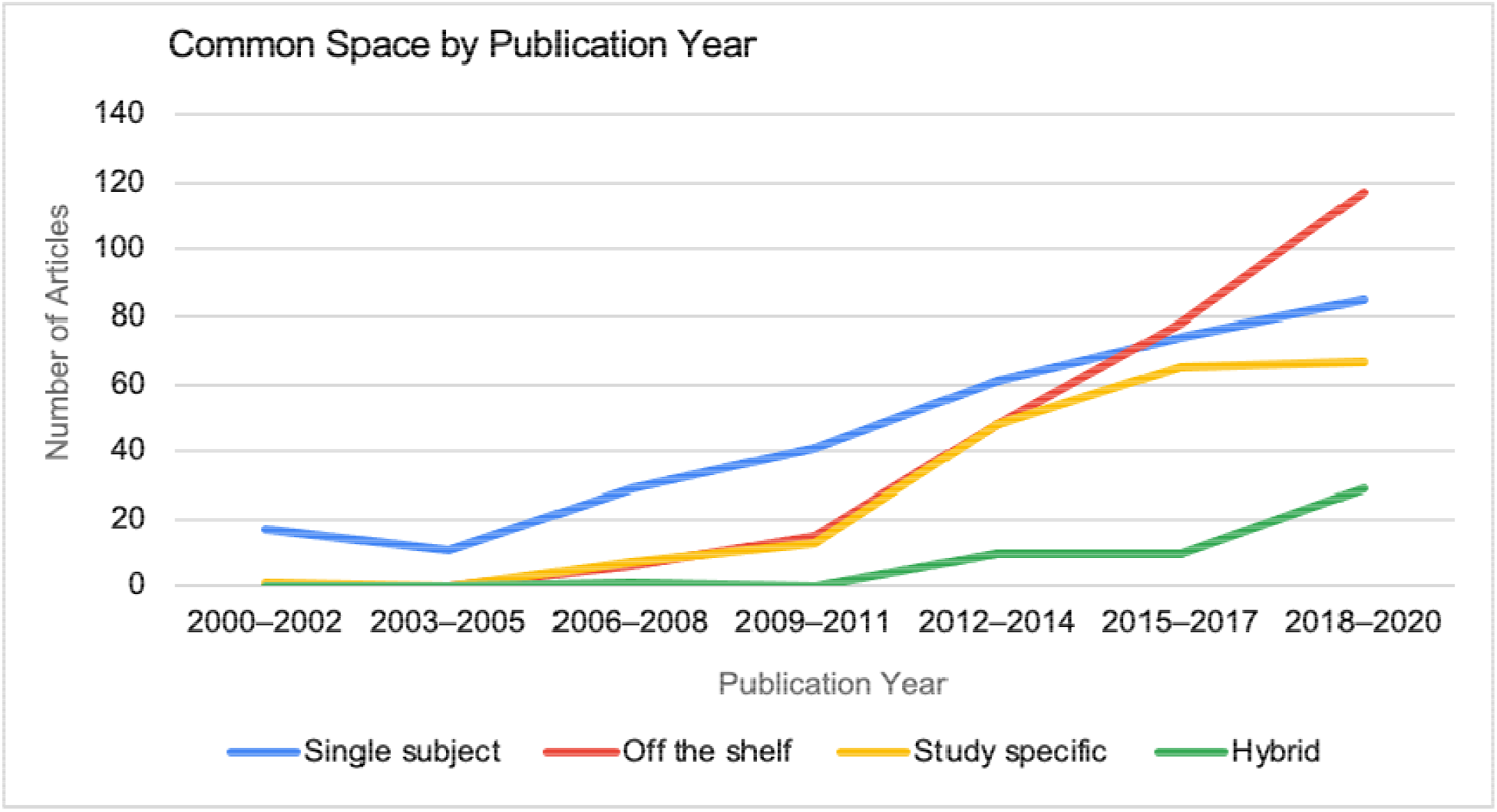
Studies published prior to 2011 primarily used single subject space compared to using a template--either predefined or study specific (single subject space: n = 98, template: n = 42). **In** contrast, studies published after 2011 primarily used a template (single subject space: n = 220, template: n = 423).

### Common space by publication year

Regarding the distribution of common space utilized by publication year, the results show a clear change in the common spaces used over time. Before 2009-2011, publications rarely used off the shelf common space. However, from 2018 to 2020, most infant MRI studies utilized an off the shelf common space. Using three 2×2 (binarized to before 2009-2011 and after 2011) contingency tables, chi-square tests suggested dependence between the distribution of common spaces by publication year was significant when comparing single subject space to off the shelf [*X^2^*(1, n = 582) = 44.96, *p* < .0001)]. The chi-square test for study specific and off the shelf (by publication year) was not significant (*p* = .44). The chi-square-test was significant for single subject space and study specific was significant [*X^2^*(1, n = 519) = 44.96, *p* < .0001)]. Additionally, we tested for a common space by publication year interaction. First, we examined the dispersion of the data. The dispersion test indicated the data was underdispersed (*α* = −1, *p* < .0001) which indicated a Poisson regression is appropriate for testing the interaction. However, the interaction terms (common space*publication year) were not significant (*ps* > .05).

### Common space by age

We examined if studies that focused on a specific age range of infants demonstrated preferences towards certain common spaces (see **Figure 5**). For this analysis we classified the studies into age groups: before term-equivalent age (TEA) referred to studies in which the sample was less than 37 weeks postmenstrual age (PMA) at the time of scan. Two additional categories were examined which included ‘longitudinal scans’ and ‘wide age range’. Longitudinal scans refer to studies that involve the same population being imaged at multiple time points. The wide age range category consisted of samples with age ranges greater than 5 months or cross-sectional data collected at multiple timepoints.

**Figure 5.**
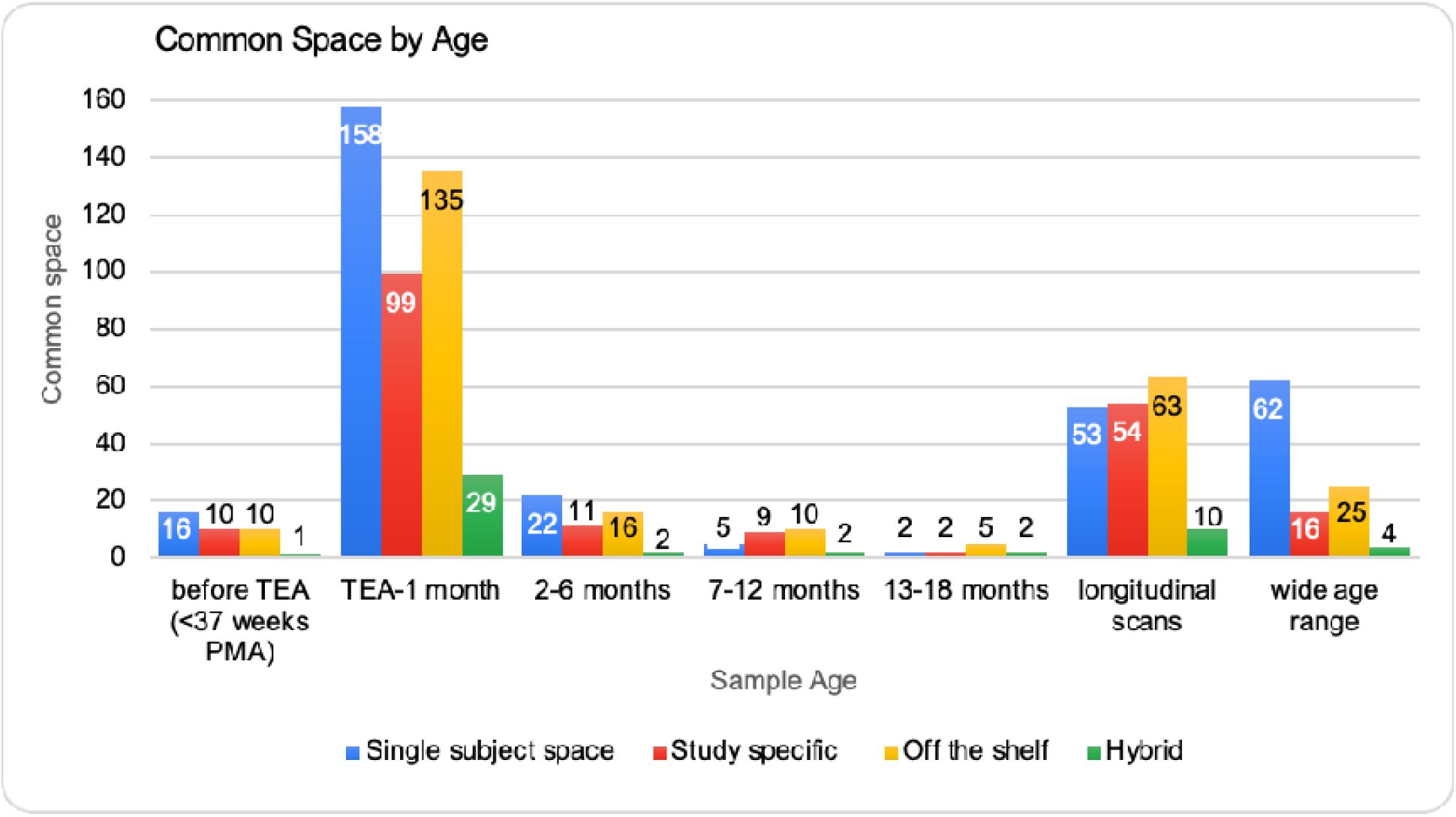
Frequency distribution of common space registrations across age groups. The distribution was analyzed in 5-month increments ranging from less than 37 weeks postmenstrual age (PMA) to 18 months. Longitudinal scans and studies utilizing a wide age range were accounted for separately.

Using three 2×2 contingency tables (publication age binarized to before TEA to 1 month old and 2-18 months old), the chi-square tests indicated for common space by age at scan, there was not significant dependence when comparing single subject space to off the shelf, study specific to off the shelf, and single subject to study specific (*ps* > .05). For both single subject space versus off the shelf (*p* < .059) and single subject versus study specific (*p* < .087), the *p*-values were at a trend level. Studies in the wide-age range group used primarily single-subject space for analyses. The use of a study specific common space stayed relatively consistent across age ranges with a slight increase for studies of 7-12-month-olds.

### Common space by special population

As infant studies may choose a specific type of common space for certain special populations of interest, we determined the number of studies using each common space for studies of preterm infants, other special populations besides preterm, and studies that did not have a special population (see **Figure 6**). The preterm category was determined by if the sample of infants imaged was less than 37 weeks gestational age at birth and included studies of low-birth-weight infants. The other special populations category included infants with autism spectrum disorder (ASD)/high risk for ASD, hypoxic-ischemic encephalopathy, prenatal exposure to alcohol/drugs/maternal mood symptoms, intrauterine growth restriction, and congenital heart diseases. These clinical categories did not have enough studies for individual group analysis.

**Figure 6.**
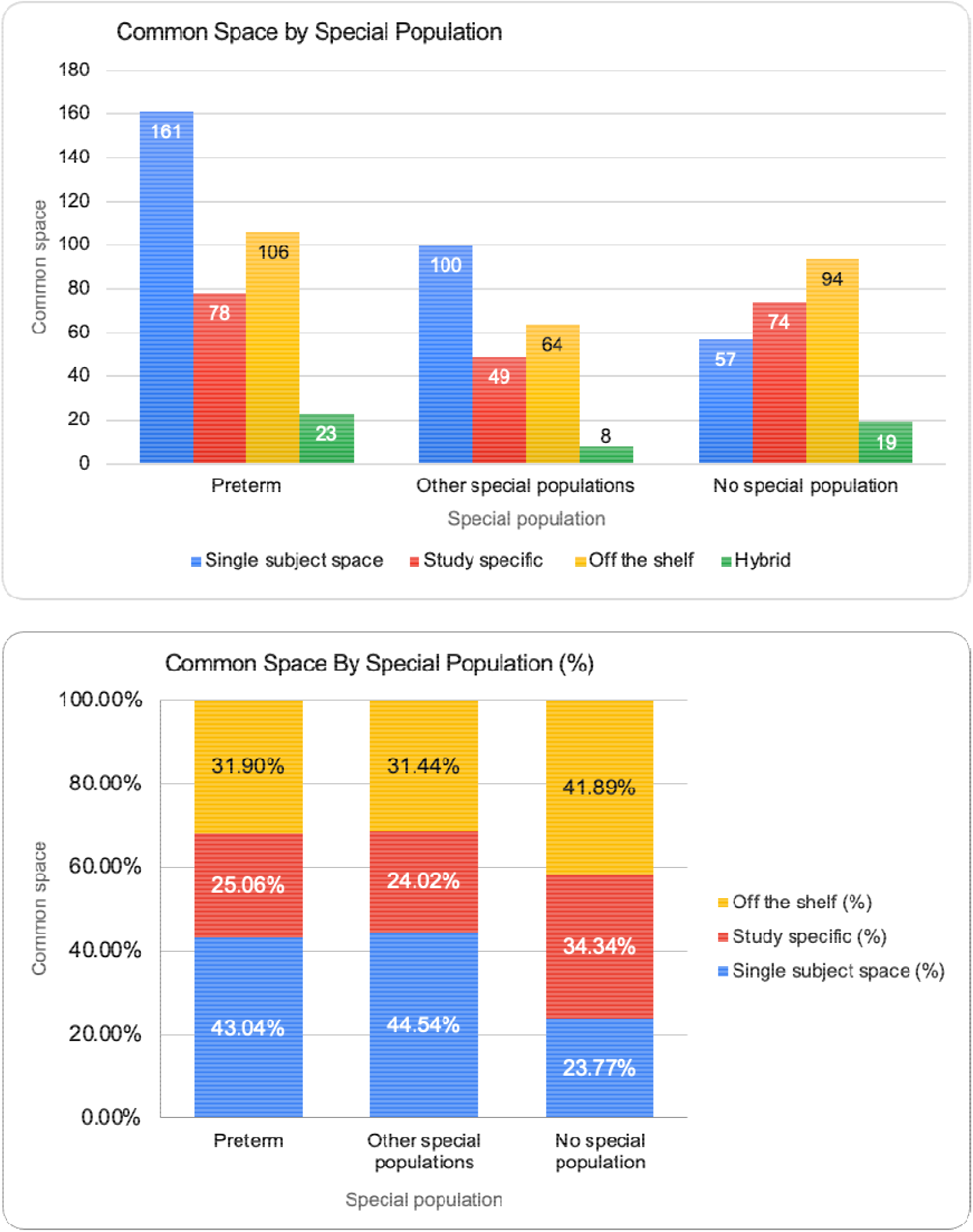
Frequency distribution (panel A) and percentage breakdown (panel B) of type of common space registration across types of population in the sample. Samples were classified as “Preterm” if they included infants who were less than 37 weeks gestational age at birth and low birth weight infants, “Other Special Populations” if they included infants with neurodevelopmental disorders, medical conditions, or prenatal exposures, or “No Special Population” if they only included typically developing infants.

The chi-square tests indicated a dependence between common space and special population for single subject space versus off the shelf [*X^2^*(2, n = 582) = 23.49, *p* < .0001)] and single subject space versus study specific common spaces [*X^2^*(2, n = 519) = 23.29, *p* < .0001)]. The chi-square test for common space by special population for study specific versus off the shelf was not significant (*p* = .95). For studies of preterm infants, single subject space was the most used method (43.04%), followed by off the shelf atlases (31.9%). For ‘other special populations’, a similar pattern was found with single subject space being most common (44.54%) followed by off the shelf atlases (31.44%). For studies that did not focus on a special population, off the shelf atlases were the most common (41.89%). The dispersion test also indicated underdispersed data (*α* = −1, *p* < .0001). The Poisson regression indicated that the interaction of common space*population was not significant (*ps* > .05).

### Off the shelf atlas distribution

Of the 287 studies classified as using an off the shelf atlas, we examined the breakdown of which atlases were the most used. The most common off the shelf atlas was the UNC infant 0-1-2 atlases (Shi et al., 2011), (http://www.med.unc.edu/bric/ideagroup/free-softwares/unc-infant-0-1-2-atlases) used in 26.5% of the studies, followed by the JHU neonate atlases (Oishi et al., 2011) (http://cmrm.med.jhmi.edu/cmrm/Data_neonate_atlas/atlas_neonate.htm) used in 16.4% of the studies, followed by 14.3% of the studies fitting into an ‘other’ category for off the shelf atlases (see **Figure 7**).

**Figure 7.**
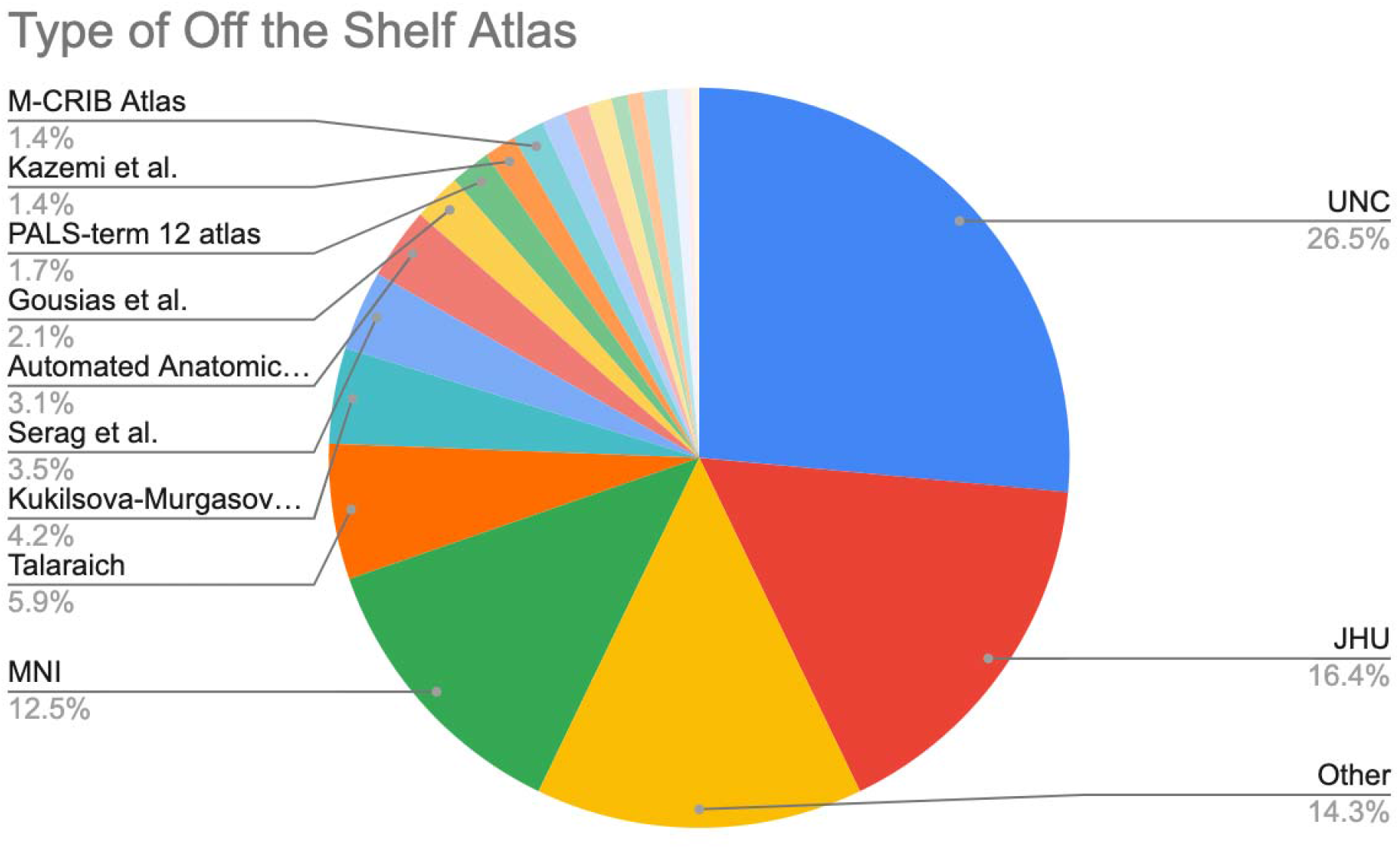
The most common predefined templates used were UNC infant atlases (n=76), JHU neonate atlases (n=47) and other; templates that were used once (n=41).

## Discussion

In our systematic review of the common spaces used for infant neuroimaging between years 2000–2020, several patterns emerged. For common spaces across modality, off the shelf atlases were used less commonly than single subject and study specific common spaces. For common spaces across publication year, DWI and sMRI studies mostly used single subject spa**ce** while fMRI studies have mostly used an off the shelf atlas. We found that single subject space was most common for neonatal studies with an increase in off the shelf atlases in older samples; off the shelf were most common for longitudinal studies (multiple scan time points across infancy). For the examination of common spaces for special populations, studies of preterm infants and other special populations favored single subject space while studies without a special population most used an off the shelf template. For studies that used an off the shelf atlas, the UNC infant atlases were the most common, followed by the JHU neonate atlases. We did not find evidence of a common space by publication year interaction or a common space by special population interaction. The findings of the systematic review indicate a need for a standardization across studies regarding a common space, that will be critical for rigor and reproducibility as the field of infant neuroimaging matures.

There still exist several challenges specific to infant neuroimaging data that have contributed to the lack of consensus and standardization of a common space across the field. A standard common space for infant neuroimaging would have to consider the rapid growth in cytoarchitecture, shape, and volume that occurs between birth and the end of the second year of life (Oishi et al., 2019). Further, compared to adult data, infant neuroimaging data has reduced tissue contrast between gray and white matter (Gilmore et al., 2018). The relative intensities of gray and white matter for T1- and T2-weighted images are similar between 4 and 8 months of age, posing issues for analyzes that involve tissue classification (Gilmore et al., 2018). In addition to poor image quality and tissue contrast, standardization of a common space for infant neuroimaging would require both a T1-weighted and T2-weighted template as researchers may only collect (or prioritize) a T1 or T2 structural image based upon the age of the sample. Lastly, the common space would have to be representative and consist of many high-quality scans. For an off the shelf common space atlases, greater than 100 images averaged is typically required (Shi et al., 2011). If this was conducted for multiple age ranges, likely, >100 high quality images acquired longitudinally would be needed. This is exceptionally difficult when imaging infants. However, as infant neuroimaging datasets increase in size, such as the Baby Connectome Project and Developing Human Connectome Project, a standard common space will be critical.

A standard common space for infant neuroimaging would need to address the existing limitations discussed above. It would also be useful for this common space to have correspondence to the adult MNI coordinate system. Most neuroimaging results are based upon adult studies (mostly registered to the adult MNI template) (Oishi et al., 2019). Therefore, having these as a reference with direct anatomical correspondence can enhance our understanding of the developing infant brain. Our results indicated a shift in the field moving from single subject space analysis to using off the shelf atlases. This finding is encouraging as both single subject space and study specific common spaces may be highly biased by the sample (S. Zhang & Arfanakis, 2013). Due to the replicability crisis (Gorgolewski & Poldrack, 2016; Klapwijk, van den Bos, Tamnes, Raschle, & Mills, 2020), it is critical for the field of infant neuroimaging to reduce as much sample-specific bias as possible to enhance rigor and reproducibility across studies. A standard common space for registration with a standard coordinate system for reporting results would facilitate meta-analyses and data sharing. For example, to directly compare the results of two studies, a research lab may need to re-register their data into the common space another lab used. This is not only time consuming but can introduce bias. Similarly, coordinate-based meta-analyses—which provide a more precise estimate of the effect size and can increase the generalizability of the results of individual studies—are not possible without a common coordinate system to report results. As infant neuroimaging has an existing small sample size issue (Korom et al., 2021), defining a standard common space will facilitate meta-analyses and data sharing and enhance rigor and reproducibility across infant neuroimaging studies.

When defining this standard common space, it is natural to ask: are currently off the shelf atlas sufficient or do new ones need to be created? The UNC infant 0-1-2 (Shi et al., 2011) and JHU neonate atlases (Oishi et al., 2011) are the most used. The UNC infant atlases are examples of spatio-temporal atlases that have a neonatal, 1 year old and 2-year-old atlas. The atlases were generated from 95 neonates that were scanned five weeks after birth and then scanned at 1 year and 2 years old. The atlases are available in both T1-weighted and T2-weighted images, tissue probability maps, and an infant Automated Anatomical Labeling (AAL) parcellation (Oishi et al., 2019; Shi et al., 2011; Tzourio-Mazoyer et al., 2002) The collection of JHU neonate atlases includes a group averaged atlas and single-subject-based atlas for T1- and T2-weighted images as well as a DTI based atlas. The group averaged atlases for the DTI and T2-weighted atlas were constructed from 20 healthy, term-birth, neonates scanned within 4 days after birth. The JHU neonate T1 atlas was constructed from 15 healthy, term-birth, neonates (37-41 gestational weeks). In terms of frequency of use, adopting the UNC infant atlases as the standard of the field could allow for future studies to be comparable to the greatest of the studies conducted to date in terms of common space. The spatio-temporal aspect of the UNC atlases is also appealing as 3 major developmental periods in early life are represented: neonatal, one years old, and two years old. However, the UNC infant atlases (0-1-2) are limited if the age of the study’s participants are outside these three time points (e.g., 5–6-month-olds, 17–18-month-olds). To mitigate this issue, spatiotemporal longitudinal 0-3-6-9-12 months-old atlases as both T1- and T2-weighted images have been released (https://www.nitrc.org/projects/infant_atlas_4d/) (Zhang et al., 2016).

Nevertheless, given the need for large sample sizes and fine grain age-specificity, these atlases may just be the starting points in developing a standard common space.

Along with the standardization of a common space (or spaces in the case of spatiotemporal atlases), the growing field of infant neuroimaging will have to adopt a standardized method of choosing an off the shelf template. For example, if two studies both have a sample of infants with a mean chronological age of 6 months and one study chooses a neonate atlas as its common space and the other chooses a 1-year atlas, the results may not be directly comparable despite the ages of the infants being similar. The choice of the different atlas for common space registration could negatively impact the rigor and reproducibility of the studies as the normalization procedure could provide differences between atlases in noise due to misregistration of the images at the voxel level and impact statistical power (Oishi et al., 2019). Standardization, not only the common space, but also how to best account for participant ages will be critical.

Infant neuroimaging datasets begin to reach “big data’’ levels like adult neuroimaging data. Two large open-source infant neuroimaging datasets exist: the Baby Connectome Project (Howell et al., 2019) and the Developing Human Connectome Project (Eyre et al., 2020). Further, the National Institutes of Health (NIH) has announced the HEALthy Brain and Child Development Study (HBCD). The HBCD will involve recruitment of a large, diverse sample of pregnant women across several sites in the United States. It will include neuroimaging data with a focus on characterizing developmental trajectories. Both within and between these large infant neuroimaging datasets consensus on common space registration (for multiple ages) will be critical to combine these valuable data.

Studies in the field of infant neuroimaging have steadily increased since the beginning of use for non-clinical studies in the early 1990s (an average of 160 publications per year to 530 per year in the last decade) (Pollatou et al., under review). However, due to the challenges of infant neuroimaging, the field has experienced a lag in standardization of best practices compared to adult MRI studies. As we have demonstrated with the current systematic review, there is a critical need for the field to establish a standard common space. To address these issues concerning establishing best practices for the field, organizations, like *Fetal, Infant, Toddler, Neuroimaging Group* (FIT’NG) (Pollatou et al., under review), will be critical to establish best practices within the field (common space, scan time, prep procedures for scanning), community exchange and collaboration (sharing analytic pipelines, datasets), and education (training across institutions at multiple levels).

The current systematic review is not without its limitations. Some of the literature currently under review lacked a clear description of methodology, thus making it hard to identify the type of registration, age range, and population. This was especially true for studies developing a study specific template. In addition, many studies used multiple common spaces or adapted an off the shelf template for their own use that was later labeled as “hybrid” or “other”. This definition did not account for the use of different common spaces for unique modalities within the same paper. Furthermore, there are some methodological limitations within our review, such as limited data on inter-rater reliability. Inclusion and exclusion inter-rater reliability were 100% when calculated in a subset of 370 papers rated by two reviewers. Clear exclusion and inclusion criteria provided guidelines and limited discrepancies between raters reviewing papers, mitigating any inter-rater reliability issues, but future reviews might benefit from collecting information about reviewer’s agreement. Finally, given the large number of papers included in our analyses, a small number of mis-classified papers is unlikely to change the general trends reported here.

## Conclusions

Despite these limitations, our systematic review provides evidence of a lack of a standard common space for infant neuroimaging studies. With the maturation of the field of infant neuroimaging, a standard common space will be critical to examine the generalizability of results across samples, ages, special populations, and imaging modalities. Further, a standard common space has the potential to increase rigor and reproducibility by reducing sample specific bias. The results of the systematic review have provided a quantification of the last two decades of infant neuroimaging to gauge where the field currently stands in terms of common space. With the results of the review in mind and an eye towards the future of the field including large consortium neuroimaging datasets, we suggest it is a critical time to adopt a standard common space.

## Acknowledgements

This publication was made possible by CTSA Grant Number TL1 TR001864 from the National Center for Advancing Translational Science (NCATS), a component of the National Institutes of Health (NIH) awarded to AJD. Its contents are solely the responsibility of the authors and do not necessarily represent the official views of the NIH.

